# Virus-Like Particles and Magnetic Microspheres Provide a Flexible and Sustainable Multiplexed Alphavirus Immunodiagnostic Platform

**DOI:** 10.1101/335315

**Authors:** Keersten M. Ricks, Charles J. Shoemaker, Lesley C. Dupuy, Olivier Flusin, Matthew A. Voorhees, Ashley N. Fulmer, Carolyn M. Six, Catherine V. Badger, Connie S. Schmaljohn, Randal J. Schoepp

## Abstract

There is a pressing need for sustainable and sensitive immunodiagnostics for use in public health efforts to understand and combat the threat of endemic and emerging infectious diseases. We describe a novel approach to immunodiagnostics based on virus-like particles (VLPs) attached to magnetic beads. This flexible, innovative immunoassay system, based on the MAGPIX® platform, improves sensitivity by up to 2-logs and has faster sample-to-answer time over traditional methods. As a proof of concept, a retroviral-based VLP, that presents the Venezuelan equine encephalitis virus E1/E2 glycoprotein antigen on its surface, was generated and coupled to magnetic beads to create VLP-conjugated microspheres (VCMs). Using these VCMs, IgG and IgM antibodies were detectable in nonhuman primate (NHP) and human clinical serum samples at dilutions of 1 × 10^4^ and greater. We extended the VCM methodology to two other New-World alphaviruses, eastern and western equine encephalitis viruses, as well as an Old-World alphavirus, Chikungunya virus, demonstrating the flexibility of this approach toward different VLP architectures. When multiplexed on the MAGPIX® platform, the VCMs provided differential diagnosis between Old-World and New-World alphaviruses and well as a route toward assessing the humoral response to both natural infection and vaccination. This VCM system will allow more rapid and efficient detection of endemic and emerging viral pathogens in human populations.

## Introduction

Immunoassays are the standard for preliminary and confirmatory diagnosis of infectious and non-infectious diseases, since they are reliable, robust, and accessible by many diagnostic laboratories. In disease outbreaks, diagnostics are the first line of defense in identification of the causative agent, treatment decisions, and eventually control and prevention of future outbreaks. An integrated diagnostic system that uses sensitive and specific molecular assays (e.g. PCR) in combination with less sensitive, but more broadly-reactive immunodiagnostic methods (e.g. ELISA) provides the highest confidence in a diagnostic result (1-3). Immunoassays are designed to detect protein-based antigens and antibodies, with antigen detection assays relying heavily on agent-specific monoclonal antibodies, and antibody detection assays requiring structurally accurate agent-specific antigens. While these immunodiagnostic technologies are relatively unsophisticated, development of sensitive and specific antibodies and antigens required for an immunoassay is often a rate-limiting step. These immunodiagnostic reagents must be sensitive, specific, robust, and most importantly, sustainable (4).

Sustainability is the most problematic issue for immunodiagnostic reagent development. A frequent viral antigen target of interest for many immunoassays is surface glycoproteins (5, 6). Traditionally, detection of anti-viral glycoprotein humoral responses for both serosurveillance and clinical diagnosis are determined by direct immunoassay methods using inactivated whole virus or lysates from infected cells to capture virus-specific antibodies in a sample. Use of whole virus for many pathogenic agents is challenging as assays must be conducted in high level biological containment (biosafety level [BSL] 3 or 4). Inactivated virus can be utilized at the BSL2 level, but vital epitopes are sometimes destroyed after inactivation protocols (7). Recently, recombinant proteins have been used as substitutes for virus preparation because they do not require specialized facilities for production, isolation, and use, and thus are more sustainable. However, soluble recombinant glycoproteins can have substantial caveats, such as the need for truncation to facilitate soluble release of the recombinant protein, artificial protein structure due to the lack of a membrane anchor or improper folding, and/or complete resistance to recombinant expression due to heterodimeric or complex maturation behavior (8-12). To circumvent these issues, an ideal diagnostic reagent would present glycoprotein antigens to the analyte material in the context of an authentic viral envelope structure while still being safe to use at the BSL2 level. In lieu of native virus, an alternate strategy for sustainable glycoprotein production can be achieved through the use of virus-like particles (VLPs). VLP generation typically relies on expression of self-assembling viral matrix proteins (e.g. HIV Gag or Ebola virus VP40) that drive particle formation via self-to-self oligomerization as well as recruitment of host cell machinery involved in vesicle formation and/or the secretory pathway (e.g. ESCRT complexes) (13). These particles are extremely safe since they contain no viral genome and can be versatile platforms for glycoprotein antigen presentation due to their well-documented ability to integrate both homologous and heterologous glycoproteins during their budding through the plasma membrane (14, 15). For analytical approaches solely dependent on reactivity, VLPs are a desirable reagent because of their ease of manufacture, antigenic fidelity, and lack of safety concerns.

Not only is development of immunoassay reagents critical when designing sustainable immunodiagnostic assays, but equally important is the choice of platform. While the traditional 96-well plate ELISA has served as a workhorse for serosurveillance efforts for decades, several immunoassay platforms have emerged to make patient sample analysis faster, more sensitive, and multiplexed at both point-of-care and centralized laboratories. One such system is the MAGPIX® developed by Luminex Corporation (Austin, Texas USA) (16). It is similar to ELISA in that it detects a typical antigen/antibody interaction, but by employing fluorescently labeled magnetic particles as a solid support, MAGPIX assays are much faster, have increased sensitivity, and better enable multiplexing (17, 18). This ability to multiplex while maintaining assay sensitivity is crucial for effective serosurveillance efforts to understand and control the spread of disease.

Herein, we outline the design and implementation of a novel diagnostic reagent, which pairs the sustainability of a VLP with the sensitivity of the MAGPIX® platform to serve as a versatile tool for detection of anti-viral glycoprotein humoral responses in a serum sample (Figure 1). More specifically, we focused on production, characterization, and optimization of alphavirus diagnostic reagents, due to previously documented problems associated with recombinant expression of their E1/E2 heterodimeric glycoproteins and because these glycoproteins play a dominant role in host immune response during alphavirus infection (19-21). As a proof of concept, Venezuelan equine encephalitis virus (VEEV) E1/E2 glycoproteins were expressed on a retroviral core VLP and conjugated to fluorescent, magnetic microspheres to create VLP-conjugated microspheres (VCMs). When incorporated onto the MAGPIX® platform, the VCMs were shown to detect both IgG and IgM in nonhuman primate (NHP) and human clinical samples with enhanced sensitivity over traditional ELISA formats in both a singleplex and multiplex format. Moreover, we employed this VCM approach to develop eastern equine encephalitis virus (EEEV), western equine encephalitis virus (WEEV), and chikungunya virus (CHIKV) assays, which when used in various multiplexed formats, provided differential diagnosis between Old-World and New-World alphaviruses, tracking of IgM response to VEEV and WEEV challenge in NHPs, and measurement of IgG response to V/E/WEEV vaccination in human serum samples. Taken together, these results demonstrate that VCMs can offer a sustainable and flexible approach for developing immunodiagnostics for use in multiple applications, including animal modeling, serosurveillance, and improved point of care diagnostics.

**Figure 1.**
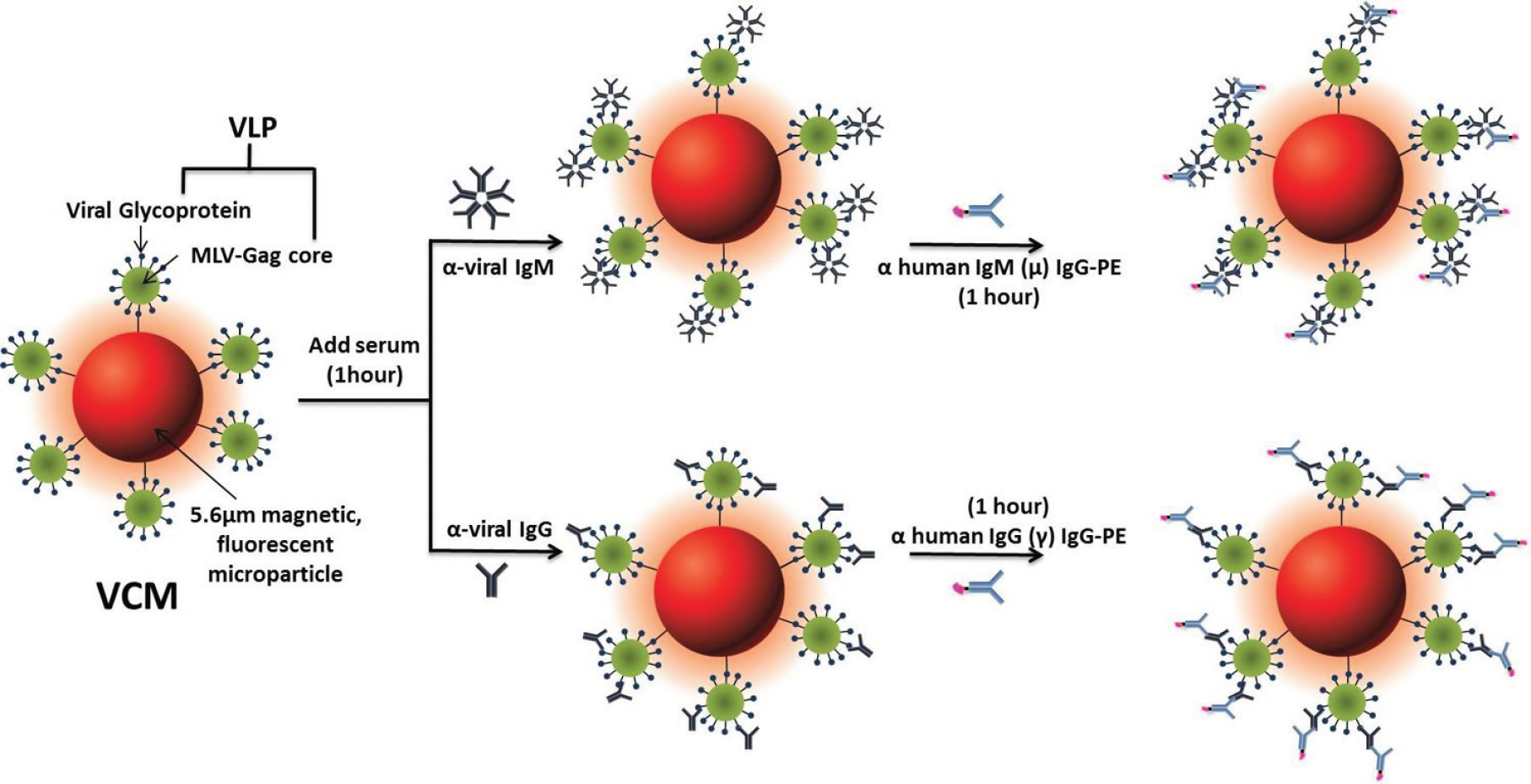
General virus-like particles (VLP) conjugated microspheres (VCM) assay schematic for detection of both antiviral IgG and IgM responses in a serum sample.

## Materials and Methods

### Antibodies and Sera

For ELISA and immuno-electron microscopy (IEM) analysis of VEEV VLPs, monoclonal antibodies (mAbs) against VEEV envelope glycoproteins E1 (3B2A9) (22) or E2 (MAB 8767, EMD Millipore, Burlington, MA, USA) were used. A mAb against VSV-G glycoprotein, (1E9F9) I14, was used as a negative control. Positive and negative control sera from NHPs vaccinated with either VEEV, EEEV, or WEEV E1/E2 plasmid DNA were generated as described previously (23). All human sera used were previously deidentified and given a “research not involving human subjects” determination. VEEV, EEEV, WEEV, and CHIKV IgG and IgM positive sera originated from either vaccination or natural infection.

### VLP Production

Plasmids encoding codon-optimized glycoprotein genes (E3-E2-6K-E1) of VEEV IAB (strain Trinidad donkey), EEEV (strain FL91-4679), and WEEV (strain CBA87) inserted into the mammalian expression pWRG7077 have been previously described (23, 24). For the construction of the Gag-encoding plasmid, the first 538 residues of murine leukemia virus (MLV) Gag-Pol ORF (GenBank: AF033811.1) were codon optimized, synthesized, and cloned into pWRG7077 using flanking 5’ *Not*I and 3’ *Bgl*II restriction sites relative to the transgene insert (Atum Inc, Menlo Park, CA, USA). HEK293T cells were seeded in T150 flasks (Corning, Inc., Corning, NY, USA) and incubated at 37°C with 5% CO2 until reaching 70-80% confluency prior to transfection with 27 μg of pWRG7077-Gag and 9 μg of pWRG7077-VEEV, pWRG7077-EEEV, or pWRG7077-WEEV E1/E2 plasmid DNA using Fugene 6 (Roche, Indianapolis, IN, USA) according to manufacturer’s instructions. Cell supernatants were collected at 24 and 48 h post-transfection, pooled, clarified by centrifugation, and filtered through a 0.45 μm filter. VLPs were concentrated through a Centricon® filter unit with a 100-kDa cutoff (EMD Millipore, Burlington, MA, USA) according to manufacturer’s instructions. VLPs were then pelleted through a 20% sucrose cushion in virus resuspension buffer (VRB; 130 mM NaCl, 20 mM HEPES, pH 7.4) by centrifugation for 2 h at 106,750 × g in an SW32 rotor at 4°C. VLP pellets were resuspended overnight in VRB at 4°C, pooled, and diluted ten-fold with VRB. The diluted VLPs were re-pelleted without a sucrose cushion as described above. VLPs were resuspended in 1/1,000 volume of VRB relative to starting supernatant and then stored at −80°C.

CHIKV VLPs were produced in a manner that has been previously described (25). Briefly, a DNA construct encoding the capsid-E3-E2-6K-E1 structural protein genes of CHIKV (strain 0706aTw) was synthesized and cloned into pWRG7077. HEK 293T cells were then transfected with 25 μg of the VLP construct using Fugene 6 as described earlier. Supernatants were harvested at 24, 48, and 72 h post-transfection and were purified as previously described. Protein concentration for all VLPs was determined by BCA assay (ThermoFisher, Waltham, MA, USA).

### Conjugation of VLPs to Magnetic Microspheres

VEEV, EEEV, WEEV, and CHIKV VLPs were conjugated to magnetic microspheres using the Luminex xMAP® antibody coupling kit (Luminex Inc., Austin, TX, USA) according to the manufacturer’s instructions. Briefly, 100 μL of Magplex microspheres (12.5 × 10^6^ microspheres/mL) were washed three times using a magnetic microcentrifuge tube holder and resuspended with 480 μL of activation buffer. Then, 10 μL of both sulfo-NHS and EDC solutions were added to the resuspended microspheres. The tube was covered with aluminum foil and placed on a benchtop rotating mixer for 20 min. After surface activation with EDC, the microspheres were washed three times with activation buffer prior to adding the VLPs at a final concentration of 10 μg VLPs/1 × 10^6^ microspheres. The tube was again covered with aluminum foil and placed on a benchtop rotating mixer for 2 h. After this coupling step, the microspheres were washed three times with wash buffer and resuspended in wash buffer to the original stock concentration of 12.5 × 10^6^ microspheres/mL for further use. The VLPs were conjugated to Magplex microsphere regions #75, #15, #45, and #25 (Luminex Inc., Austin, TX, USA), respectively, in order to facilitate multiplexing experiments.

### Detection of anti-viral IgG or IgM in NHP or Human Sera using VLP-coupled Magplex Microspheres

VCMs were diluted 1:250 in phosphate buffer saline (PBS) with 0.02% Tween-20 (PBST) and added to the wells of a Costar polystyrene 96-well plate at 50 μL per well (2500 microspheres/well). The plate was placed on a Luminex plate magnet, covered with foil, and microspheres were allowed to collect for 60 sec. While still attached to the magnet, the buffer was removed from the plate by shaking. Then, 50 μL of serum, diluted in PBST with 5% skim milk (PBST-SK) was added to appropriate wells and the plate was covered and incubated with shaking for 1 h at room temperature (RT). The plate was washed three times with 100 μL of PBST using the plate magnet to retain the Magplex microspheres in the wells and then 50 μL of a 1:100 dilution of goat anti-human IgG (H&L) phycoerythrin conjugate (Sigma-Aldrich, St. Louis, MO, USA) or goat anti-human IgM (anti-mu) phycoerythrin conjugate (Abcam, Cambridge, UK) in PBST-SK were added to the wells. The plate was covered and incubated with shaking for 1 h at RT. After incubation, the plate was washed three times and the Magplex microspheres were resuspended in 100 μL of PBST for analysis on the MAGPIX®. For multiplexed experiments, the VLP-coupled microspheres were each added to the plate so that 2500 microspheres of each VCM set were dispensed per well. The remainder of the multiplexed assay was performed as described above. For sera from VEEV, EEEV, and WEEV-infected NHPs, analysis was conducted in a BSL-3 suite with infected material handled in a Class II Biological Safety Cabinet. All samples were run in triplicate.

## Results

### Synthesis and Characterization of VLPs

MLV-based VLPs were chosen for VEEV glycoprotein presentation because they are high yielding, homogenous, and can accommodate a wide range of glycoprotein antigens (26, 27). Transient expression of two DNA plasmids encoding both VEEV E1/E2 and the first 538 amino acids of MLV Gag in mammalian cells generated highly homogenous particles presenting both the E1 and E2 VEEV glycoproteins on their surface as determined by electron microscopy (Figure 2A, B). A molar ratio of 3:1 Gag plasmid to VEEV E1/E2 plasmid yielded the highest incorporation of the glycoproteins into the particles (SI Figure 1B). This same ratio was optimal for the EEEV and WEEV E1/E2 glycoproteins (SI Figure 1C) as well as other non-alphaviral glycoproteins that have been tested (data not shown). Comparison of the VEEV E1/E2 VLPs against γ-irradiated, whole TC-83 VEEV antigen was made by direct ELISA. Plates coated with equal amounts of each antigen were probed with either E1 or E2-specific mAbs or with sera from NHPs vaccinated with the VEEV E1/E2 plasmid DNA (Figure 2 C, D). The VLPs performed better as compared to the inactivated material with respect to binding of the mAbs and displayed equivalent reactivity to polyclonal sera from vaccinated NHPs. As further proof that the MLV-based VEEV VLP glycoproteins were present in a native, functional conformation, we demonstrated successful entry of the VLPs into target cells. This entry was also blocked by neutralizing polyclonal sera from vaccinated NHPs, further supporting the native-like structure of the VLP-embedded glycoproteins (SI Figure 1A). VLPs bearing glycoproteins for two other encephalitic alphaviruses, EEEV and WEEV, were generated in an analogous fashion. They were characterized by both western blot (SI Figure 1C) and ELISA (data not shown). CHIKV VLPs were developed using an alternate VLP architecture that is dependent on the ability of CHIKV capsid and envelope proteins to spontaneously drive VLP formation (28). Relying on this organic approach, CHIKV VLPs were generated and characterized by both western blot (SI Figure 1C) and ELISA (data not shown).

**Figure 2.**
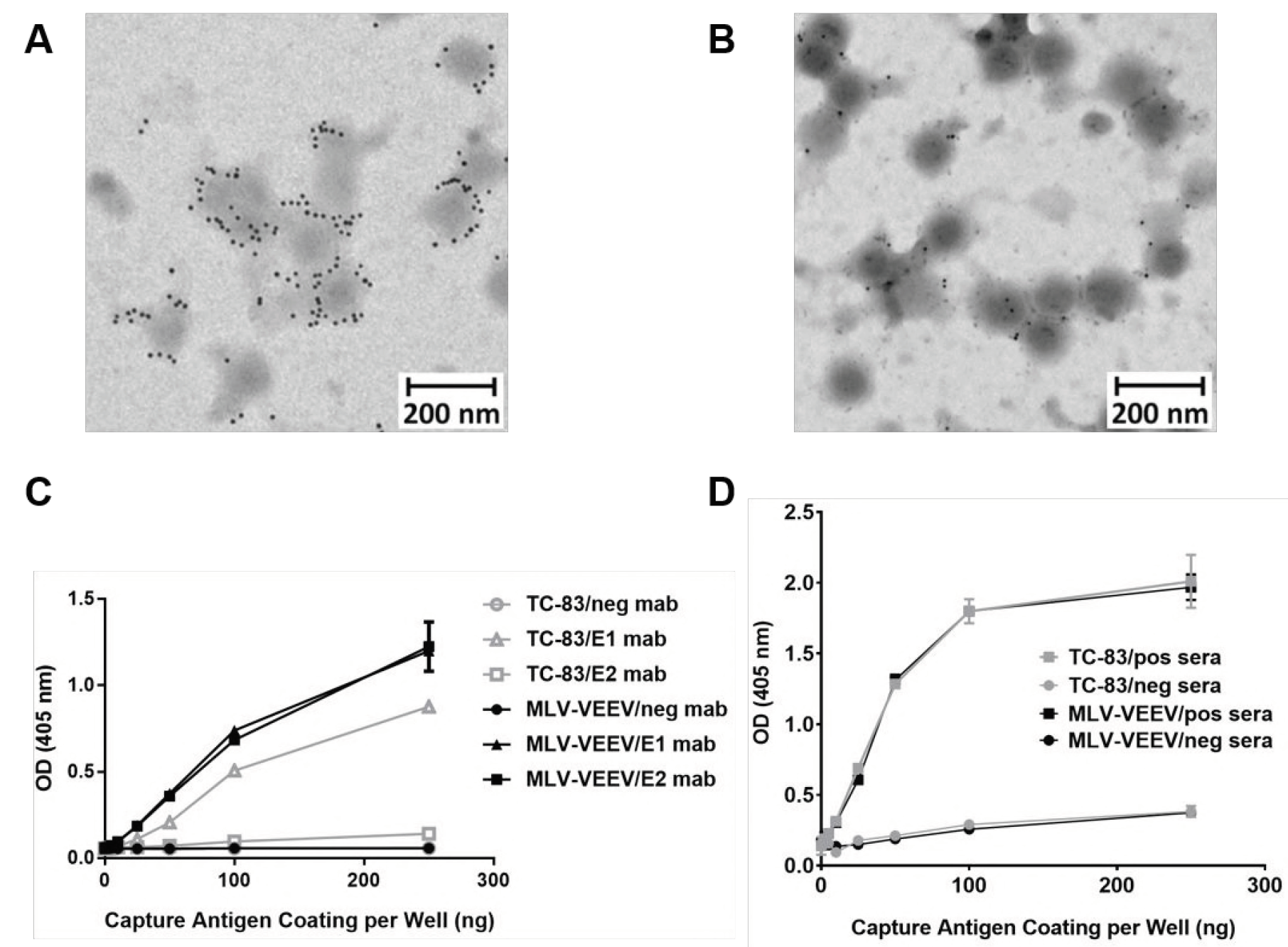
Immunoelectron microscopy (IEM) imaging of VEEV VLPs stained with (**A**) anti-E1 or (**B**) anti-E2 mAbs. VLPs were imaged at 80,000× magnification. Scale bars are indicated for 200 nm. (**C**) VEEV VLPs or γ-irradiated TC-83 virus were coated onto ELISA plates at indicated amounts and probed with mAbs against either VEEV E1 or E2 or a negative control mouse antibody or (**D**) negative and VEEV-reactive sera from NHPs. Error bars represent standard deviation.

### Conjugation, Characterization, and Comparison to Traditional ELISA of VEEV VCMs

VEEV VLPs were conjugated to Magplex microspheres using carbodiimide coupling chemistry to covalently link the amine groups from the surface glycoproteins of the VLP to the carboxylate surface of the microparticle. Saturation of the particle surface with VLPs was observed at a concentration of 10 μg/million microspheres, so this was chosen as the standard loading concentration for the VCMs (SI Figure 1D). Upon screening known anti-VEEV IgG positive NHP sera with the VCMs on the MAGPIX®, the limit of detection (LoD) was determined to be at a 1 × 10^5^ dilution in assay buffer (Figure 3A). Signal at this dilution was significantly higher when compared to the same dilution of negative NHP sera (t-test; p<0.0001). The VEEV VCM MAGPIX® assay was two orders of magnitude more sensitive toward IgG detection than traditional 96-well ELISA assays using inactivated TC-83 cell lysate or VEEV VLP direct capture antigens (Figure 3B).

**Figure 3.**
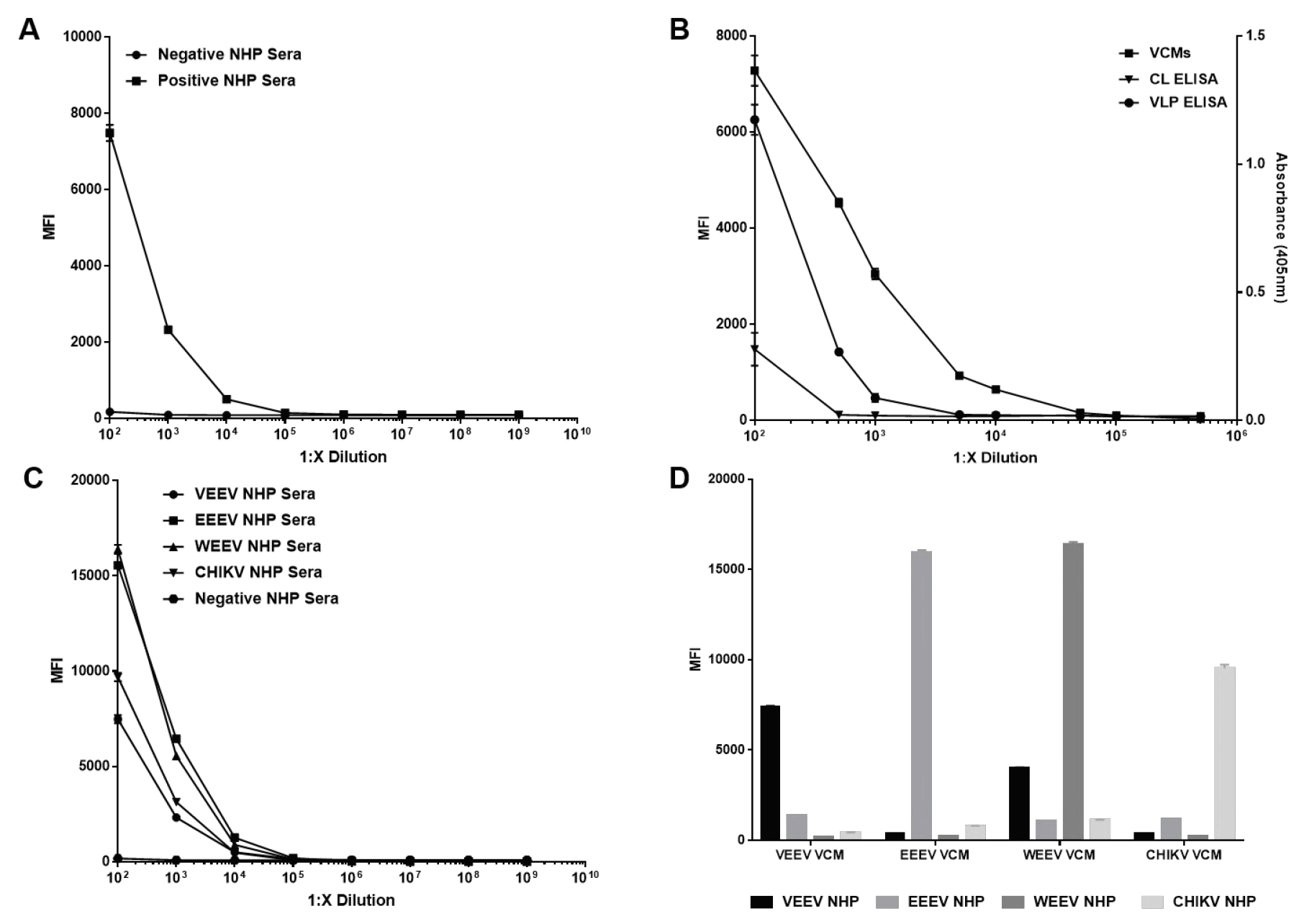
(A) Limit of detection (LoD) of anti-VEEV IgG in nonhuman primate (NHP) sera using the VEEV VCM MAGPIX assay. Signal was statistically significant (p=0.01; unpaired t-test) over baseline to a dilution of 1 × 10^5^, (B) Comparison of VEEV VCM assay to traditional direct ELISA based on VEEV infected cell lysate (CL) or VEEV VLP as capture antigen for detection of anti-VEEV IgG in NHP sera. (C) LoD for each alphavirus VCM assay with corresponding positive NHP sera. LoD for all four singleplex assays fell at a dilution of 1 x 10^5^, (D) VEEV, EEEV, WEEV, and CHIKV multiplex VCM assays for detection of anti-alphavirus IgG in positive NHP sera at a 1:100 dilution. Error bars represent standard deviation.

### Characterization of Alphavirus VCMs in Singleplex and Multiplex Formats

Given the success of coupling and testing the VEEV VCMs, the three other alphavirus VLPs, namely EEEV, WEEV, and CHIKV, were coupled to Magplex beads at a loading concentration of 10 μg/million beads and screened against known IgG positive NHP sera in a singleplex format (Figure 3C). The limit of detection of each singleplex alphavirus VCM assay was similar to that observed in the proof of concept test with the VEEV VCMs at a 1 × 10^5^ serum dilution. While LoD was not affected by multiplexing the alphavirus VCMs and interrogating individual NHP sera (SI Figure 2), it was necessary to determine whether the assay would be specific for the etiologic agent or whether crossreactivity would be observed (19). VEEV, EEEV, WEEV, and CHIKV NHP sera were screened for IgG using a mixture of the four alphavirus VCMs (Figure 3D). At a 1:100 dilution of each NHP sera, some crossreactivity was observed as signal was significantly above baseline for several of the non-correlative VCM-NHP sera pairings, most notably between the VEEV NHP sera and the WEEV VCM; however, much of this crossreactivity diminished at a 1:1,000 dilution of sera in assay buffer (SI Figure 3).

### Diagnostic Utility of Alphavirus VCMs for Animal Models and Human Clinical Samples

While the VCMs proved to be highly sensitive for detection of IgG in convalescent NHP sera, the diagnostic utility of such a platform lies in its sensitivity toward IgM detection in sera from both animal models and human clinical samples at early time points post-infection. The presence of pathogen-specific IgM represents the earliest antibody response of an organism to infection. As the course of infection progresses toward convalescence, the presence of IgM generally decreases as IgG rises to dominate the humoral response (29). VEEV, EEEV, and WEEV NHP sera (n=4 for each cohort) at multiple time points post-challenge were screened using a triplex V/E/WEEV VCM assay. Anti-VEEV and anti-WEEV IgM response over baseline was observed at days 5, 7 and 9 post-infection (Figure 4). Interestingly, no anti-EEEV IgM response was observed at any time point (data not shown). This was potentially due to the EEEV aerosol challenge model used, which resulted in 75% lethality by day 5 post-exposure and 100% by day 7. Some IgM crossreactivity was observed when the time points were screened in the triplex assay, most notably that of the EEEV VCMs with the VEEV NHP sera (SI Figure 4).

**Figure 4.**
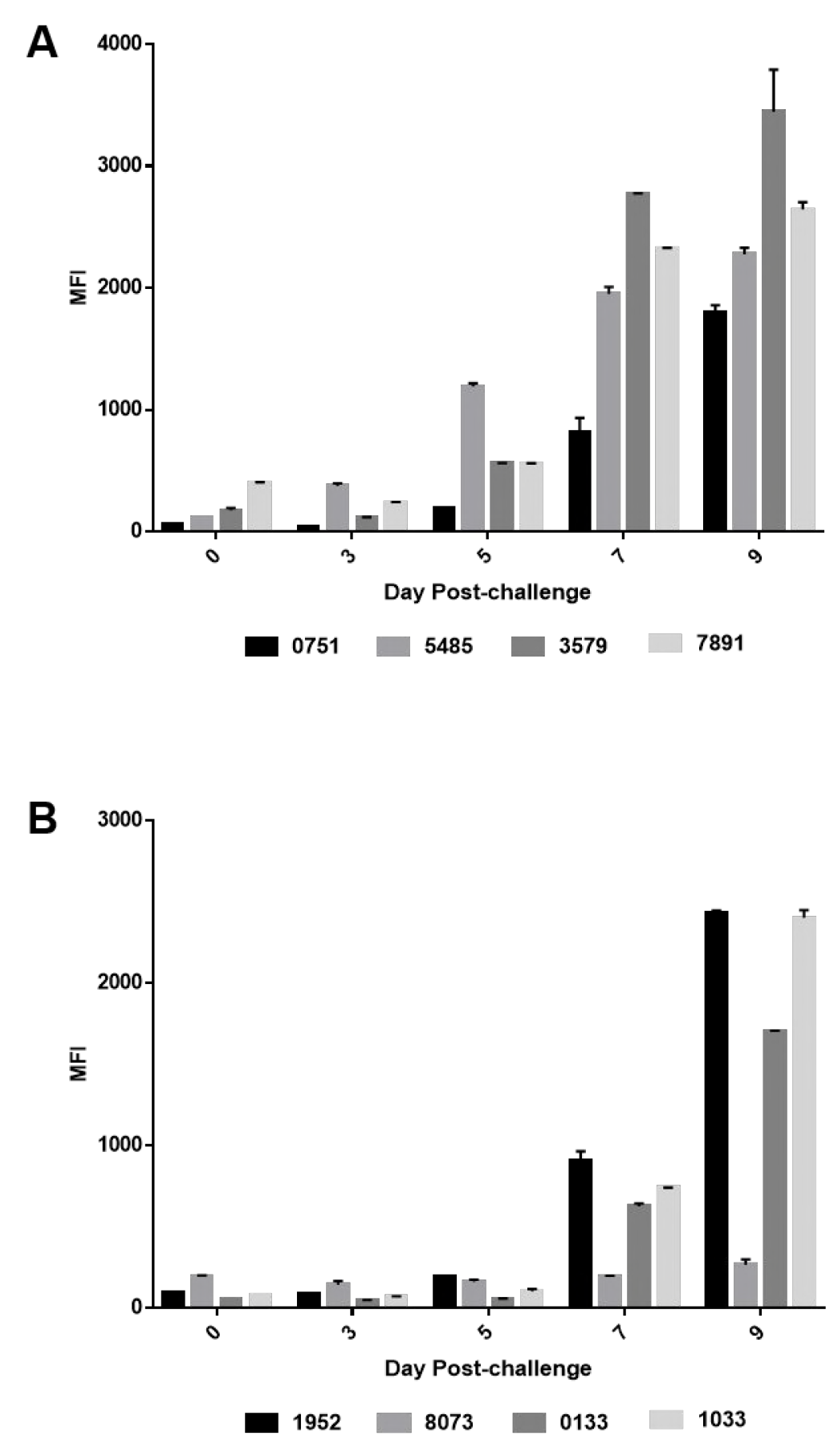
Detection of (A) anti-VEEV IgM and (B) anti-WEEV IgM in VEEV and WEEV challenged NHPs, respectively, at Days 0, 3, 5, 7, and 9 post-challenge. All NHP sera was diluted 1:100 in assay buffer.

Clinical samples from human vaccinations were screened with the same V/E/WEEV triplex VCM assay for the presence of anti-viral-IgG. Thirty-two total samples were screened, where 16 were known negatives, a cohort of 10 were vaccinated with VEEV (TC-83), EEEV, and WEEV vaccines, and a cohort of 6 with VEEV IND vaccine only (Figure 5). The VEEV vaccine used was live-attenuated TC-83 whereas the EEEV and WEEV vaccines were both formalin-inactivated. The V/E/WEEV VCM triplex had a sensitivity and specificity of 100%, with 0% false positive and false negative rates, for detection of anti-V/E/WEEV IgG in the triple vaccine cohort. For the cohort of 6 VEEV only vaccinations, the V/E/WEEV triplex assay had a sensitivity and specificity of 100% toward VEEV IgG detection, but a false positive rate of 6% and 11% for the EEEV and WEEV assays, respectively.

**Figure 5.**
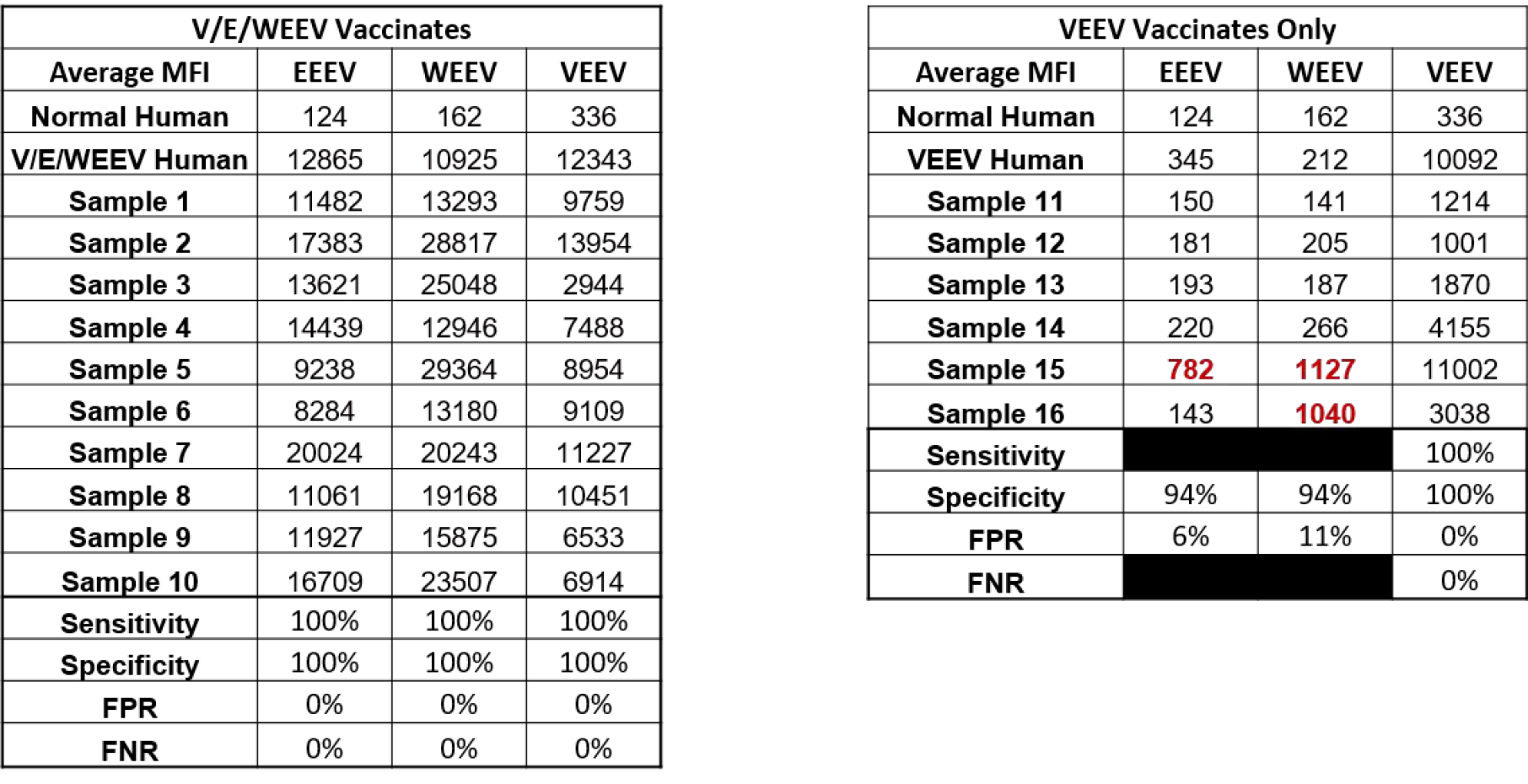
IgG response to V/E/WEEV or VEEV only vaccination using a triplex V/E/WEEV VCM assay. A cohort of 10 participants received a V/E/WEEV trivalent vaccine while a cohort of 6 received VEEV-only. 16 known negative samples were also tested by the triplex assay. Sensitivity was calculated as True Positives (TP)/(True Positives (TP) + False Negatives (FN)). Specificity was calculated as True Negatives (TN)/(True Negatives (TN) + False Positives (FP)). False positive rate (FPR) was calculated as FP/(FP+TN) and false negative rate (FNR) as FN/(TP+FN). The values highlighted in red indicate false positive values from the VEEV-only cohort in the triplex assay. All samples were diluted 1:100 in assay buffer.

To further explore alphavirus crossreactivity, human clinical sera was screened using a VEEV/CHIKV duplex VCM assay for the presence of anti-IgG and IgM antibodies. As singleplex assays, the VEEV and CHIKV VCMs are highly sensitive toward IgG and IgM detection in correlating human sera (SI Figure 5). LoDs at a dilution of 1 x 10^5^ and 1 x 10^4^ were observed for VEEV IgG and IgM, respectively. Likewise, LoDs for CHIKV IgG and IgM fell at dilutions of 1 x 10^6^ and 1 x 10^4^, respectively. When these same IgG and IgM positive sera were screened in a duplex VEEV/CHIKV assay at a 1:100 dilution, there was clear specificity of the IgG response from the CHIKV sera toward the corresponding VCM, with minimal crossreactivity with the VEEV VCM (Figure 6A). No crossreactivity of the IgG response from the VEEV sera was observed with the CHIKV VCM. When IgM positive sera was screened in the duplex assay, no crossreactivity was observed (Figure 6B).

**Figure 6.**
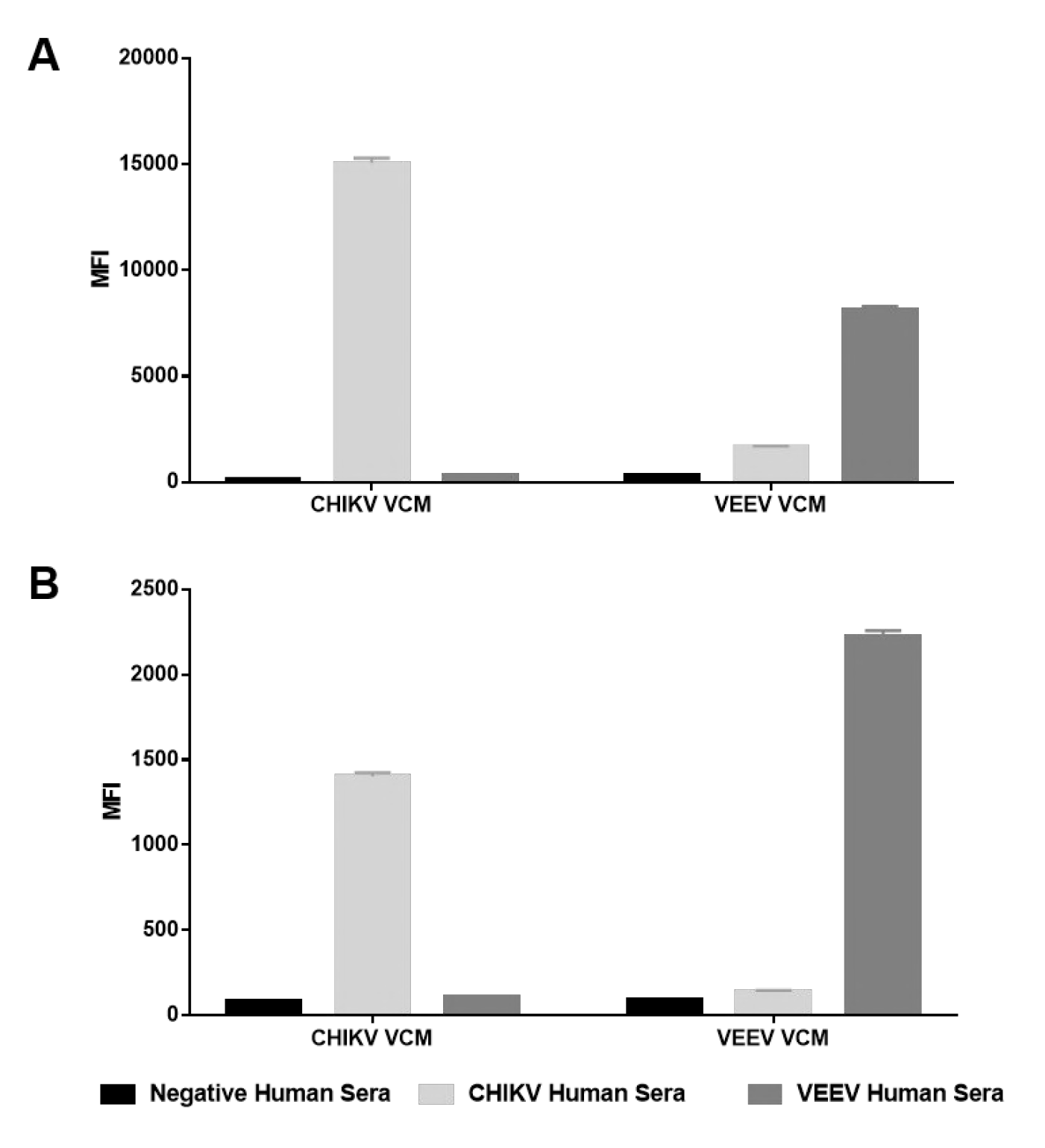
Discrimination between Old-World and New-World alphaviruses for (A) IgG and (B) IgM detection from human clinical sera at a 1:100 dilution.

## Discussion

In developing countries, access to sensitive and sustainable diagnostics at healthcare facilities can be a key determinant between controlling isolated cases of emerging or re-emerging infectious disease or progression to an outbreak (12, 30). Enveloped viruses cause the vast majority of pathogenic viral diseases affecting human populations. Glycoproteins on the surface of these viruses typically elicit robust antibody responses and therefore often represent the best target antigen for detecting the serological response to these infectious agents. We developed a multiplexed immunodiagnostic assay for detecting antibody response to these glycoproteins in the context of alphavirus infections. Alphaviruses, which are members of the *Togaviridae* family, are of concern both as emerging, enzootic pathogens and as biothreat agents (31). While VEEV is a NIAID priority B pathogen and an area of concern for biodefense agencies, CHIKV is a major global health concern, as its widespread prevalence leads to severe disease burden and long-term disability (19). We designed a novel immunodiagnostic reagent for detection of anti-glycoprotein antibody responses to VEEV infection and demonstrated how the methodology can be extended to additional alphaviruses such as EEEV, WEEV, and CHIKV. In an effort to meet the key criteria needed for effective serological surveillance and clinical diagnostic protocols for these pathogens and other biothreats, we aimed to rationally design the VCM reagent to be safe, sensitive, flexible, and, most importantly, sustainable when used in field-forward platforms. While many traditional immunoassays utilize inactivated viral preparations or recombinant antigens as capture reagents, these reagents can be costly to produce, lack structural fidelity to native particle-associated antigen, and, in the case of inactivated BSL3/4 agents, require high level biocontainment for production and inactivation. Using VLPs as capture antigens is highly advantageous, since they are easy and inexpensive to produce, inherently safe, and maintain native structural antigen conformation. Integrating the surface glycoproteins onto either a heterologous retroviral particle core or a homologous core (i.e. CHIKV) was a rational approach for creating diagnostically useful, membrane-stabilized glycoprotein targets for immunodiagnostic applications. As our results demonstrated, these particles were simple to produce and reactive against seropositive NHP sera and glycoprotein-specific mAbs (Figure 2).

The VCM as an antibody target combined with the MAGPIX® to detect antibodies to multiple targets simultaneously in a single sample is an extremely flexible platform technology. VEEV NHP sera, screened with VEEV VCMs, yielded an IgG LoD at serum dilution of 1 × 10^5^ (Figure 3A). A marked improvement over traditional ELISA was also observed with the VCM approach, not only in a reduction in time-to-answer, but with a two-order magnitude of increased IgG sensitivity (Figure 3B). As an extension of this concept, three other alphavirus VCMs were developed and characterized—each yielding IgG LoDs at serum dilutions 1 × 10^5^ for corresponding NHP serum (Figure 3C). When assays were multiplexed for VEEV, EEEV, WEEV, and CHIKV antibody detection, there was no observed loss in sensitivity with the combined assay components (SI Figure 2). Some anti-glycoprotein IgG crossreactivity amongst members of the New-World alphaviruses was observed when sera were screened with the quadruplex assay; however, this was to be expected as the glycoproteins, in particular E1, of some alphavirus family members share regions of homology and possess conserved epitopes (32). IgM responses in NHPs to VEEV and WEEV challenge were observed starting at day 5 and day 7 post-challenge, respectively (Figure 4). Minimal crossreactivity with the VEEV and EEEV VCMs was observed with the WEEV sera, but significant crossreactivity was observed on the EEEV VCM with the VEEV sera. Nonetheless, it was apparent there was a strong correlation between the VCM with the highest signal response and the etiologic agent of the screened sera. Pairing this multiplexed VCM immunoassay with PCR, in an integrated diagnostic approach, would lead to the highest confidence in specific pathogen identification.

Human sera from equine encephalitic virus vaccines were screened alongside a cohort of true negatives with a triplex V/E/WEEV VCM assay to determine assay performance metrics for IgG detection (Figure 5). For the cohort of 10 vaccinations that received a trivalent V/E/WEEV vaccine, the assay had a sensitivity and specificity of 100% with 0% false positive and false negative rates. This was to be expected as it was known that this cohort received the trivalent vaccine; however the assay served to confirm seroconversion. For the cohort of 6 vaccinations that received only the VEEV vaccine, the triplex assay had a sensitivity and specificity of 100% for the VEEV assay, but the false positive rates for the EEEV and WEEV assay components were 6% and 11%, respectively. This observation is likely attributed to the previously noted crossreactivity within IgG positive equine encephalitic sera (33, 34). Despite this crossreactivity, there was again strong overall signal on the VEEV VCM, correlating to the etiologic agent of the vaccine. When human clinical sera from VEEV and CHIKV infection were screened in a duplex with VEEV and CHIKV VCMs, less crossreactivity was observed, specifically with IgM detection (Figure 6). This would suggest that differential diagnosis by IgM detection is achievable for identifying Old-World versus New-World alphaviruses, which would be significant in regions where both the equine encephalitic viruses and CHIKV are endemic.

A sustainable, flexible, and robust immunoassay such as the VCM platform presented here utilized on a field-forward platform such as the MAGPIX® has the potential to greatly improve diagnostic capacity for both diagnostic and serosurveillance scenarios, as this platform affords higher throughput, faster read times, reduced sample consumption, and increased sensitivity to reach a diagnostic result as compared to screening by traditional ELISA methods. Many overseas centralized laboratories are currently able to perform basic ELISA immunoassays, but are often limited to commercially available kits. These kits are costly, time consuming to run, variable in performance from lot to lot, and difficult to keep in stock, as manufacturers often discontinue products or have long purchasing lead times. We were able to screen multiple serum sample types using several multiplexed assay iterations in two hr on the same plate using only 2 μL of each sample, whereas up to 24 hr, separate plates, and up to 10 μL of each sample would be required to achieve the same result with a traditional 96-well plate ELISA. While we demonstrated the ability of our VCM platform to detect of alphavirus immune responses in animal models, as well as human clinical samples (both natural infection and vaccinated), this VCM platform technology could be extended to other virus families, particularly those endemic in central and western sub-Saharan Africa, to create regional-specific serosurveillance or diagnostic panels for use at centralized testing facilities. In particular, the VCM approach will be useful for diagnostic strategies targeting antibody response against other arbovirus envelope targets (e.g. dengue virus E, etc.), whose glycoproteins are frequently difficult to produce in recombinant form (35). Incorporation of sustainable and sensitive immunoassay reagents such as VCMs into field-forward technologies will become increasingly important in global surveillance and diagnostic efforts to limit the spread of infectious viral diseases.

## Acknowledgements

The authors would like to thank Dr. Jay Hooper and Dr. Steve Kwilas for providing the αVSV-G mAb (1E9F9) I14 and Dr. Judith White and Dr. James Simmons for the generous gift of the beta-lactamase/Gag fusion protein. We also would like to thank Cindy Rossi, Scott Olschner, Brian Kearney, Tammy Clements, and Dr. Mark Poli for their helpful discussions. KMR and CJS were funded by the National Research Council Research Associateship Award at US Army Medical Research Institute of Infectious Diseases (USAMRIID), as well as support from Oak Ridge National Laboratories. The laboratory work was funded in part by the Armed Forces Health Surveillance Branch (AFHSB) and its Global Emerging Infections Surveillance (GEIS) Section through USAMRIID, ProMIS projects P0012_18_RD and P0017_17_RD.

Opinions, interpretations, conclusions, and recommendations are those of the authors and are not necessarily endorsed by the U.S. Army.

Research on human subjects was conducted in compliance with DoD, Federal, and State statutes and regulations relating to the protection of human subjects, and adheres to principles identified in the Belmont Report (1979). All data and human subjects research were previously deidentified and given a “research not involving human subjects” determination by the USAMRIID Office of Human Use and Ethics, NHSR Determination FY17-26. The V/E/WEEV vaccine study was conducted under IRB approved protocols FY05-23 and HP05-23.

Research on NHPs was conducted under an IACUC approved protocol, V05-17, in compliance with the Animal Welfare Act, PHS Policy, and other federal statutes and regulations relating to animals and experiments involving animals. The facility where this research was conducted is accredited by the Association for Assessment and Accreditation of Laboratory Animal Care, International and adheres to principles stated in the Guide for the Care and Use of Laboratory Animals, National Research Council, 2011.

